# Designing Minimal Genomes Using Whole-Cell Models

**DOI:** 10.1101/344564

**Authors:** Joshua Rees, Oliver Chalkley, Sophie Landon, Oliver Purcell, Lucia Marucci, Claire Grierson

## Abstract

In the future, entire genomes tailored to specific functions and environments could be designed using computational tools. However, computational tools for genome design are currently scarce. Here we present algorithms that enable the use of design-simulate-test cycles for genome design, using genome minimisation as a proof-of-concept. Minimal genomes are ideal for this purpose as they have a simple functional assay, the cell either replicates or not. We used the first (and currently only published) whole-cell model, for the bacterium *Mycoplasma genitalium*. Our computational design-simulate-test cycles discovered novel *in-silico* minimal genomes smaller than *JCVI-Syn3.0*, a bacteria with, currently, the smallest genome that can be grown in pure culture. In the process, we identified 10 low essentiality genes, 18 high essentiality genes, and produced evidence for at least two *Mycoplasma genitalium in-silico* minimal genomes. This work brings combined computational and laboratory genome engineering a step closer.

## Introduction

For genome-scale engineering and design, minimal genomes are currently the best proof-of-concept ^1^. These are reduced genomes containing only genes essential for life, provided there is a rich growth medium and no external stressors ^1,2^. The largest scale efforts in genome minimisation to date include: *JCVI-Syn3.0*, a 50% gene reduction of *Mycoplasma mycoides* ^2^; *several strains of Escherichia coli* reduced by 38.9% ^3^ and 35% ^4^ of their base pairs *in-vivo*; an *E.coli* gene reduction of 77.6% in *Saccharomyces cerevisiae* ^5^; *and two 36% gene reductions of Bacillus subtilis* ^6^. Initially, these were either prescriptively designed, with requirements based on current biological knowledge, or based on extensive laboratory testing of individual genes. These were then developed iteratively in the lab, a time consuming and expensive process due to the limitations of current techniques and unexpected cell death, likely caused by unknown genetic interactions. This hinders progress as laboratories can only follow a small number of high-risk research avenues with limited ability to backtrack ^1^.

Another approach, building novel organisms from the bottom-up, is currently infeasible in the majority of bacteria due to technological and economic constraints. Megabase sized genomes can be constructed within yeast ^5,7^, but one of the most promising approaches, genome transplantation, has only been demonstrated in a subset of *Mycoplasmas* ^8–10^ and is mutagenic ^9^.

A further barrier to genome minimisation is the dynamic nature of gene essentiality. A simple definition of a cell as “living” is if it can reproduce, an “essential” gene being indispensable for cell division. A *“*non-essential*”* gene can be removed and leave division intact ^1,11^. But a cell’s need for specific genes (and their products) is dependent on the external cellular environment and on the genomic context ^1^ (the presence or absence of other genes, and resulting gene products, in the genome), which can change each time a gene is removed. Some essential genes can become dispensable with the removal of a particular gene (i.e. a toxic byproduct is no longer produced, so its removal is unnecessary), referred to as “protective essential” genes ^1,12,13^. Likewise, some non-essential genes become essential when a functionally equivalent gene is removed, leaving a single pathway to a metabolite (a “redundant essential” gene pair). Additionally, gene products can perform together as a complex, with individually non-essential genes involved in producing an essential function ^14^; when enough deletions accumulate to disrupt the group, the remaining genes become essential. The cellular death that occurs when redundant essential genes are removed together, or complexes are disrupted, is referred to as synthetic lethality ^2,15,16^. A recent review ^1^ updates gene essentiality from a binary categorisation to a gradient with four categories: no essentiality (if dispensable in all contexts), low essentiality (if dispensable in some contexts, i.e. redundant essential and complexes), high essentiality (if indispensable in most contexts, i.e. protective essential), and complete essentiality (if indispensable in all contexts). These broad labels describe an individual gene’s essentiality in different genomic contexts, and are compatible with other labels that explain underlying mechanisms and interactions in greater levels of detail.

To overcome the above, large-scale problems we used existing computational models with novel genome design algorithms to investigate 10,000s of gene knockout combinations *in-silico*, with rapid feedback and iteration. Testing potential genome reductions at scale for lethal interactions should produce functional *in-silico* genomes, which can be implemented *in-vivo* with a lower risk of failure. This generation of non-prescriptive designs, with no assumed biological requirements outside those inherent in the model, increases the likelihood of novel findings.

We used the *Mycoplasma genitalium* (*M.genitalium*) whole-cell model ^17^, which describes the smallest culturable, self-replicating, natural organism ^18^ (at the time the model was built). It is the only existing model of a cell’s individual molecules that includes the function of every known gene product (401 of the 525 *M.genitalium* genes), making it capable of modelling genes in their genomic context ^17^. A single cell is simulated from random initial conditions until the cell divides or reaches a time limit. The model combines 28 cellular submodels, with parameters from >900 publications and >1,900 experimental observations, resulting in 79% accuracy for single-gene knockout essentiality ^17^. Outside of single-gene knockout simulations, it has been used to investigate discrepancies between the model and real-world measurements ^17,19^, design synthetic genetic circuits in the context of the cell ^20^, and make predictions about the use of existing antibiotics against new targets ^21^.

We produced two genome design algorithms (Minesweeper and the Guess/Add/Mate Algorithm (GAMA)) which use the *M.genitalium* whole-cell model to generate minimal genome designs. Using these computational tools we found functional *in-silico* minimal genomes, between 33 and 53 genes smaller than the most recent predictions for a reduced *Mycoplasma* genome of 413 genes ^2,15,16^. These *in-silico* genomes are ideal candidates for further *in-vivo* testing.

## Results

### Genome Design Tools: Minesweeper and GAMA

Minesweeper and GAMA conduct whole-cell model simulations in three step cycles: design (algorithms select possible gene deletions); simulate (the genome minus those deletions); and test (analyse the *in-silico* cell produced). Simulations that produce dividing cells go through to the next cycle of simulations. The number of gene deletions increases in each cycle, producing progressively smaller genomes. Minesweeper and GAMA have generated 2157 and 53,451 of *in-silico* genomes respectively to date, but for brevity only the smallest genomes are presented here.

Minesweeper is a four stage algorithm inspired by divide and conquer algorithms ^22^, initially investigating genes individually to identify complete/high essentiality genes, before breaking the genome into differently sized subsets to broadly test, then accumulating deletions and identifying low essential genes as they appear. It deletes genes in groups that get progressively smaller until it reaches individual gene deletions, and only deletes non-essential genes (as determined by single-gene knockout simulations, see Initial Input below). By not considering essential genes the search area is reduced, which makes it capable of producing minimal genome size reductions quickly (within two days). It uses between 8 and 359 CPUs depending on the stage, with data storage handled by user submitted information and simulation execution conducted manually.

GAMA is a biased genetic algorithm ^23^. It first conducts two stages (Guess and Add) of only non-essential gene deletions, which form a biased initial generation for the next (Mate) stage. The latter follows a standard genetic algorithm process. GAMA produces deletion segments that vary by individual genes, requiring 100s-1000s of CPUs. It takes two months to generate minimal genome size reductions as it uses between 400 and 3000 CPUs depending on the stage. Custom management code is used to coordinate and execute simulations, and store data.

### Initial Input

To generate an initial input for Minesweeper and GAMA we simulated single-gene knockouts in an otherwise unmodified *M.genitalium in-silico* genome (as previously reported ^17,19^, Supplementary Information A). The 359 protein-coding genes were simulated individually (10 replicates each), with 152 genes being classified as non-essential and 207 genes classified as essential (i.e. producing a dividing or nondividing *in-silico* cell, respectively). The majority of genes (58%) are essential; this was expected, as *Mycoplasmas* are obligate parasites with reduced genetic redundancy ^24^.

318 genes showed consistent results across knockout replicates, the same phenotype in 10/10 cases, with 41 showing inconsistent results. Statistical analysis (binomial proportion confidence interval, Pearson-Klopper, 95% CIs for: one 6/10 replicate [5.74, 6.87], 7/10 replicates [6.66, 7.93], 8/10 replicates [7.56, 8.97], 9/10 replicates[8.45, 9.99]) resulted in the genes being classified by the majority phenotype (see Methods and Supplementary Information B & N). Overall, our results agree 97% with Karr et.al ^17^, see Supplementary Information C.

### Minesweeper Method and Results

The first stage of Minesweeper conducts individual gene knockouts *in-silico* to identify complete/high essentiality genes, removing them as gene deletion candidates.

The second stage sorts the remaining non-essential genes into deletion segments (from 12.5 to 100% of the remaining genes (Figure 1) resulting in 26 segments, broadly sweeping for potential low essential genes. The deletion segments that produce a dividing *in-silico* cell are carried forward to the next stage.

**Figure 1.**
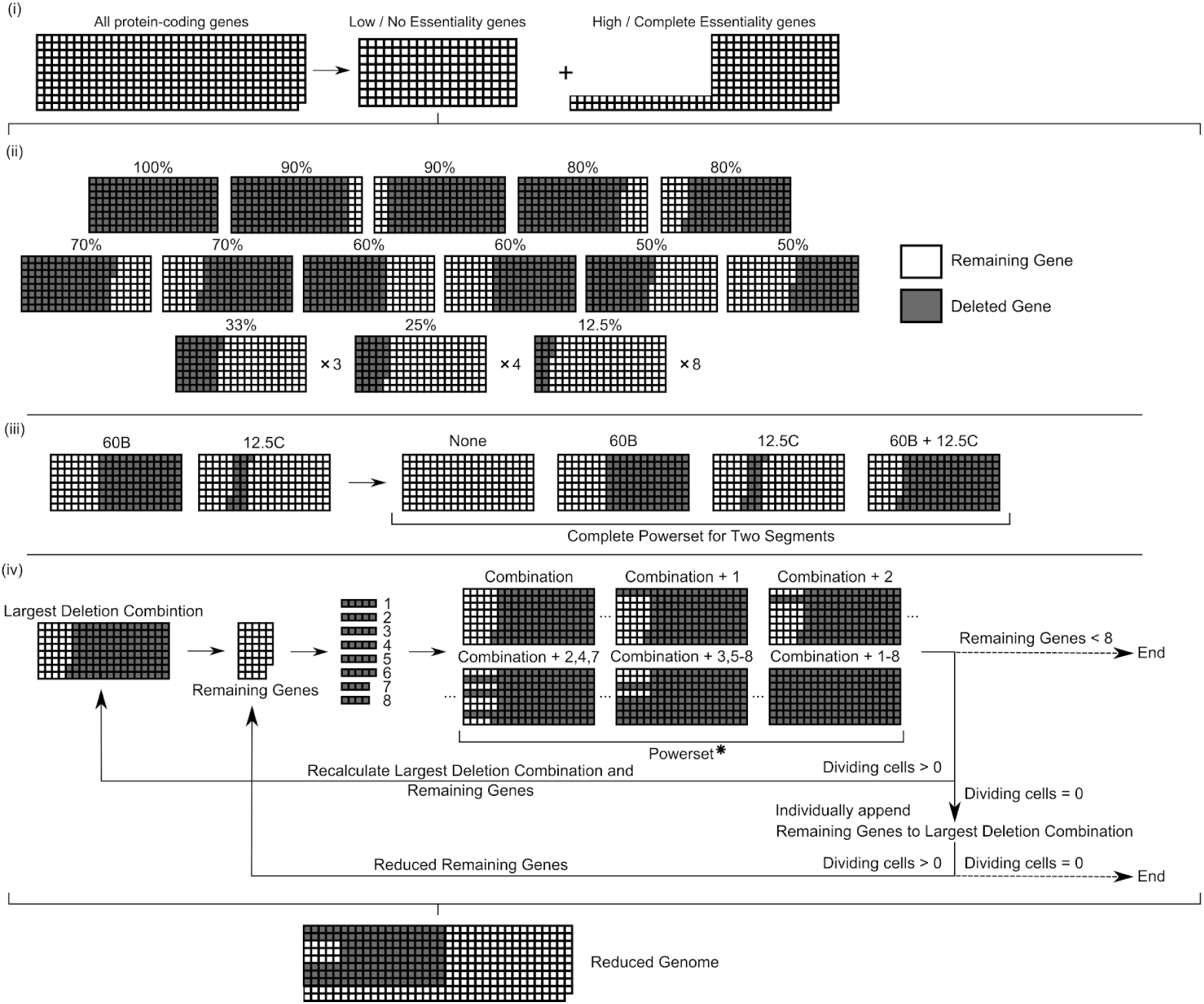
Minesweeper Algorithm for Genome Design. (i) *in-silico* single gene knockouts are conducted to identify low / no essentiality genes (whose knockout does not prevent cell division). (ii) 26 deletion segments, ranging in size from 100% to 12.5% of the low / no essentiality genes, are simulated. Grey indicates a gene deletion, white indicates a remaining gene. Deletion segments that do not prevent division go to the next stage. (iii) The largest deletion segment is matched with all dividing, non-overlapping segments. A powerset (all possible unique combinations of this set of matched deletion segments) is generated and each combination simulated. Deletion segments that do not prevent division go to the next stage. (iv) The largest deletion segment determines the remaining low / no essentiality genes that have been deleted. These remaining genes are divided into eight groups (see Methods), a powerset generated for these eight groups, and each member of the powerset individually appended to the current largest deletion combination and simulated. If none of these simulations produces a dividing cell, the remaining genes are appended as single knockouts to the current largest deletion combination and simulated. The individual remaining genes that don’t produce a dividing cell are temporarily excluded and a reduced remaining gene list produced. Details of simulations settings are available in the Methods. Powerset* = the complete powerset is not displayed here.

The third stage progresses with the largest deletion segment that produced a dividing cell, which is matched with other dividing, non-overlapping segments. A powerset (all possible unique combinations of the matched segments) is generated, and each combination of deletion segments is simulated in an *in-silico* cell.

The fourth stage is cyclical. The largest deletion combination that produces a dividing cell is used to generate a remaining gene list, those yet to be deleted, which narrows down potential conditional essential genes. It splits the remaining genes into eight groups (see Methods) and a powerset is generated. Each combination is individually appended to the current largest deletion combination and simulated. Again, the largest deletion combination that produces a dividing cell is used to generate a remaining gene list, which is used to start the next cycle of the stage.

If none of the combinations produces a dividing cell, the remaining genes are singly appended to the largest deletion combination and simulated. The individual remaining genes that don’t produce a dividing cell are temporarily excluded and a reduced remaining gene list is produced, which is used at the start of the next cycle.

The fourth stage continues until there are eight or less remaining genes (where a final appended powerset is run) or all individually appended remaining genes do not produce a dividing cell. Both outcomes result in a list of deleted genes and identified low essential genes.

Minesweeper produced results quickly, within two days the third stage removed 123 genes (a 34% reduction) comparable to current lab-based efforts in other species ^3,4,6^. The repeating fourth stage increased the overall number of deletions.

In total, Minesweeper deleted 145 genes (Figure 1), creating an *in-silico M.genitalium* cell containing 256 genes (named Minesweeper_256), which replicates DNA, produces RNA and protein, grows, and divides.

### GAMA Method and Results

The first and second stages of GAMA (Guess and Add) are pre-processing stages that provide input for the third stage (Mate), a genetic algorithm. Typically a genetic algorithm would start with random gene knockouts, but to reduce the number of generations required to produce minimal genome size reductions, the Mate stage starts with large gene knockouts produced by Guess and Add (Figure 2).

**Figure 2.**
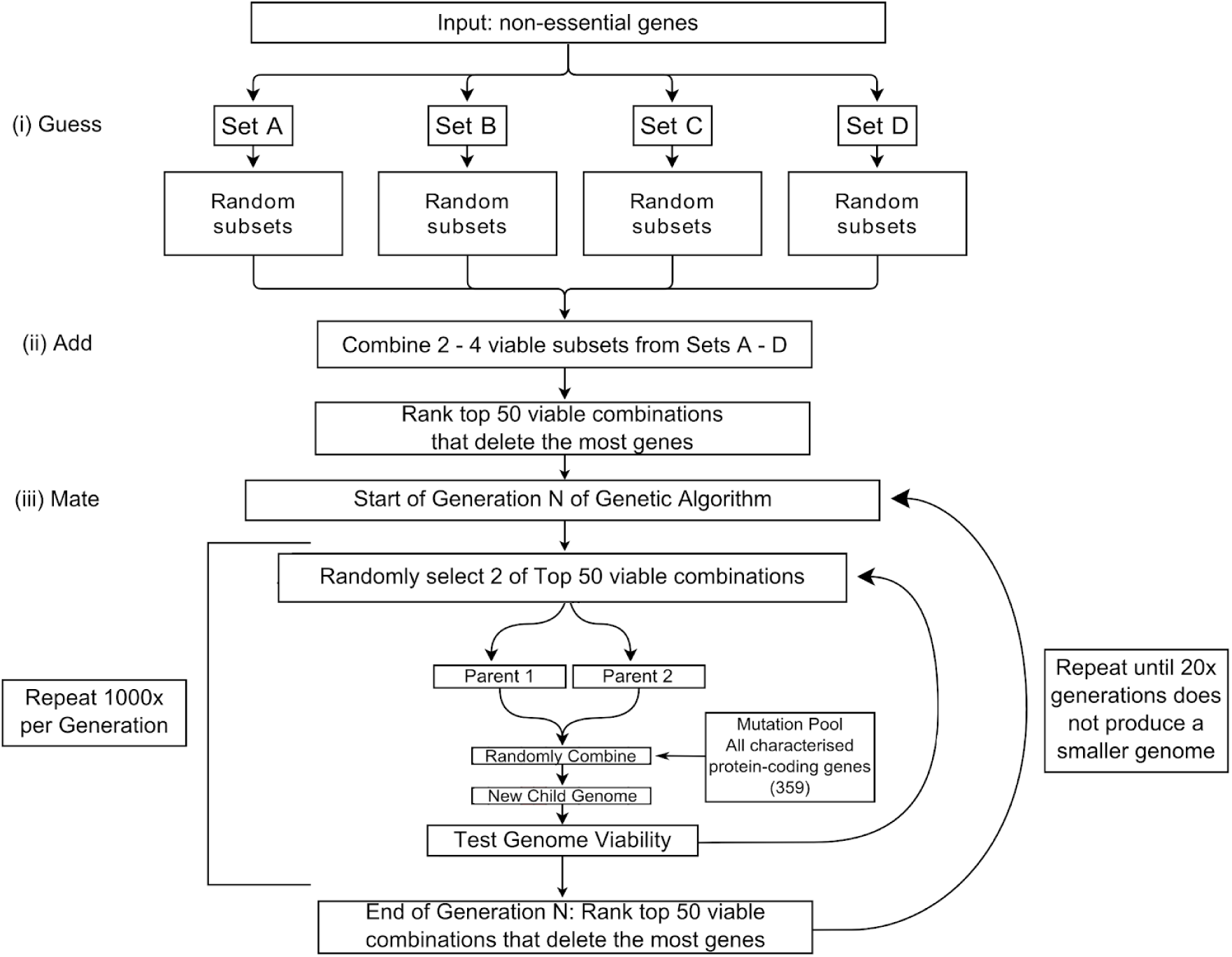
GAMA Algorithm for Genome Design. (i) Only non-essential genes whose knockout does not prevent cell division are deletion candidates and are equally divided into Sets A - D. 400 random subsets are produced and simulated per set, each containing 50-100% of the genes within the set. Deletion segments that do not prevent division (“viable”) go to the next stage. (ii) 3000 combinations are generated and simulated. (iii) Is a cyclical step. The mutation pool targets a random number of genes for alteration (both knockins and knockouts), including essential genes. Details of simulations settings are available in the Methods.

In the first stage, Guess, all the non-essential genes from the initial input are segmented into four sets, to reduce the size and number of combinations to search through. Each set is then used to generate ∼400 subsets, by randomly choosing combinations of 50 - 100% of the genes (∼40) in the set to delete. The build and test steps are then conducted. If a cell divides, the deletion subset is labelled “viable” and carried forward to the next stage.

During the second stage, Add, a number of “viable” subsets are randomly selected from two, three or four of the sets, which are combined into a larger set. Being able to select smaller numbers of subsets reduces the chance of only producing non-dividing cells. ∼3000 combined subsets are created, simulated and tested. Those producing a dividing cell are ranked based on the number of genes deleted. The 50 smallest genomes are taken forward to the mate stage.

During the third stage, Mate, the 50 smallest genomes are used to speed up the discovery of minimal genomes. The mate stage is cyclical, consisting of generations containing 1000 simulations. Each simulation in a generation combines two of the 50 smallest *in-silico* genomes at random, and introduces random gene knockouts and knock-ins from a pool of all protein-coding genes (including complete and high essentiality genes). The genomes produced are ranked and compared to the smallest 50 genomes, with the new smallest 50 being carried through to the next generation. The mate step automatically stops after 100 generations, but was manually stopped at 46 generations, after 20 generations without producing a smaller genome.

In total, the smallest GAMA-reduced *in-silico* genome deleted 165 genes, creating an *in-silico M.genitalium* genome of 236 genes (named GAMA_236). GAMA removed more genes than the Minesweeper method, while still producing a simulated cell which replicates DNA, produces RNA and protein, grows, and divides.

### GAMA_236 and Minesweeper_256 Genomes

We investigated the characteristics of our two minimal genomes in terms of how consistently they produced a dividing *in-silico* cell, and the range of possible behaviour they displayed. We simulated 100 replicates of an unmodified *M.genitalium in-silico* genome, Minesweeper_256, GAMA_236, and a single-gene knockout of a known essential gene (MG_006) to provide a comparison (see Supplementary Information G). The rate of division (or not in the MG_006 knockout simulations) was analysed to assign a phenotype penetrance percentage, quantifying how often an expected phenotype occurred. The unmodified *M.genitalium* and MG_006 knockout *in-silico* genomes demonstrated consistent phenotypes (99% and 0% divided, respectively). Minesweeper_256 was slightly less consistent (89% divided), while GAMA_236 was substantially less consistent, producing a dividing *in-silico* cell 18% of the time. This is not entirely unexpected given the greater number of gene deletions affecting essential gene functions (according to the GO term analysis).

The 100 replicates for the unmodified *M.genitalium* genome, Minesweeper_256, and GAMA_236 were plotted to assess the range of behaviour (Figure 3). The unmodified *M.genitalium* whole-cell model (Figure 3, top row) shows the range of expected behaviour for a dividing cell (in line with previous results ^17^). Growth, protein production, and cellular mass increase over time, with most cells dividing at around 10 hours, though division can occur between 6 and 11 hours. RNA production fluctuates but increases over time. DNA replication follows a characteristic shape, with some simulations delaying the initiation of DNA replication past ∼9 hours.

**Figure 3.**
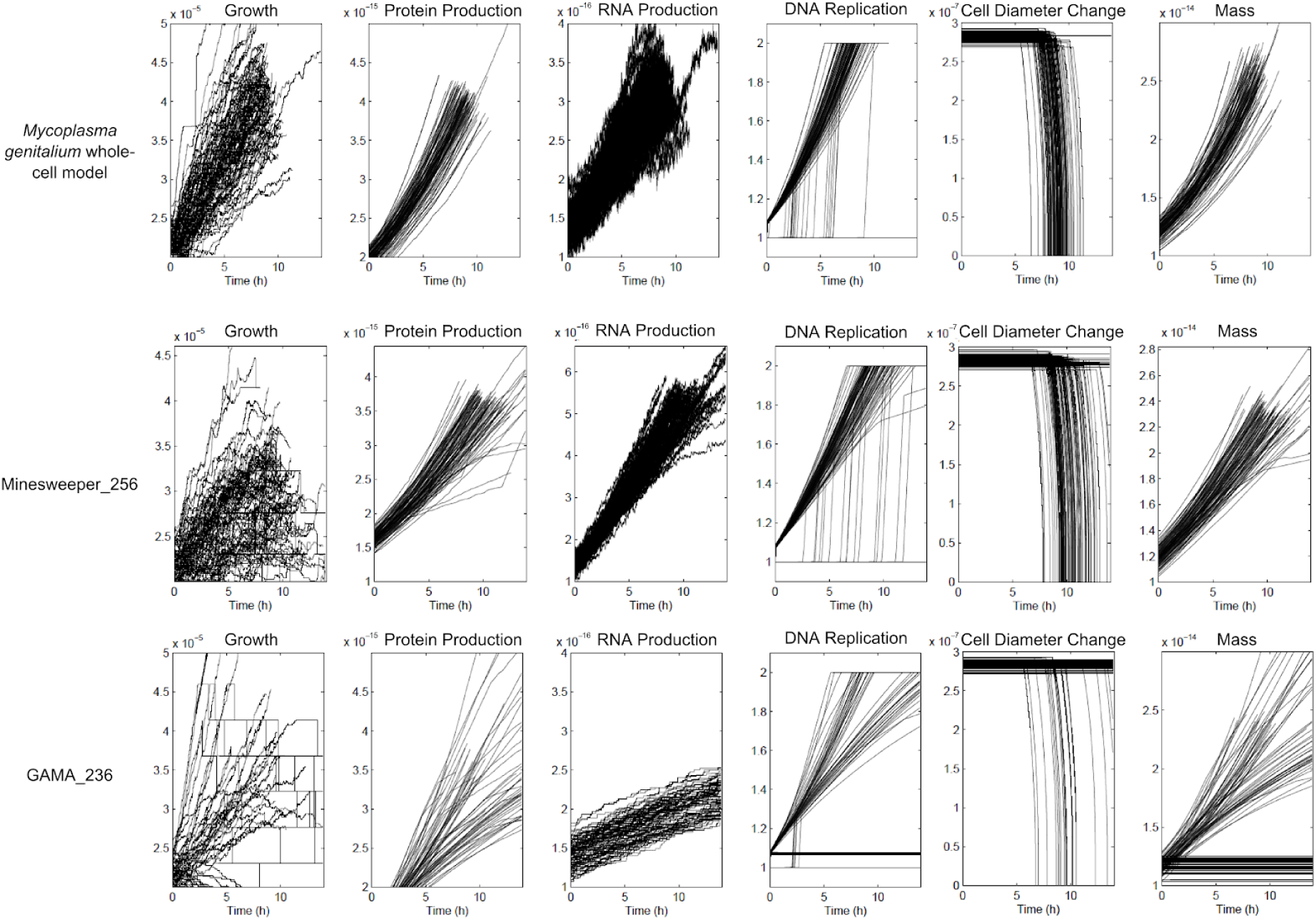
Comparison of unmodified *Mycoplasma genitalium* whole-cell model, Minesweeper_256, and GAMA_236 outputs. 100 *in-silico* replicates, with second-by-second values plotted for 6 cellular variables over 13.89 hours (the default endtime of the simulations). Top row is unmodified genome, showing the expected cellular behaviour (previously show by Karr et al ^17^) and is used for comparison. Minesweeper_256 and GAMA_236 show deviations in phenotype caused by gene deletions. Non aggregated data for each *in-silico* simulation is available (see Methods).

By comparison, Minesweeper_256 (Figure 3, middle row) displays slower, and in some cases decreasing, growth over time which is capped to a lower maximum. Protein production and cellular mass are generated more slowly and present some erratic behaviour. The range of RNA production is narrower compared to the unmodified *M.genitalium* whole-cell model. DNA replication takes longer and initiation can occur later (at 11 hours). Cell division occurs later, between 8 and 13.889 hours. A number of simulations can be seen failing to replicate DNA and divide.

Compared to the other genomes, GAMA_236 (Figure 3, bottom row) shows a much greater range of growth rates. Some grow as fast as the unmodified genome, some are comparable to Minesweeper_256, and some show very low or decreasing growth. Observable protein levels appear between 2 and 5 hours, followed by a slower rate of protein production in some simulations. Cellular mass is either similar to Minesweeper_256 or slower. The range of RNA production is reduced and the rate of RNA production is slower.

Some simulations replicate DNA at a rate comparable to the unmodified genome, others replicate more slowly, and some do not complete DNA replication. Cell division occurs across a greater range (6 - 13.889 hours). A number of simulations showing metabolic defects can be seen. These do not produce any growth, and can also be seen failing to replicate DNA and divide.

We investigated what processes were removed in the creation of Minesweeper_256, using gene ontology (GO) biological process terms (see Methods and Supplementary Information I-K). The baseline *M.genitalium* whole-cell model has 259 genes of 401 genes (72% coverage) with GO terms on UniProt ^25^. Minesweeper_256 has 186 (73%) genes with GO terms and 70 (27%) genes without. The 140 gene deletions reduced 22 (14%) GO categories, and removed 41 (27%) GO categories entirely, of which 29 (70%) were associated with a single gene (see Supplementary Information L).

The GO categories reduced include: DNA (replication, topological change, transcription regulation and initiation); protein (folding and transport); RNA processing; creation of lipids; cell cycle; and cell division. As the *in-silico* cells continue to function, we can assume that these categories could withstand low-level disruption.

Removed GO categories that involved multiple genes include: proton transport; host interaction; DNA recombination and repair; protein secretion and targeting to membrane; and response to oxidative stress.

Removed GO categories that contain single genes include: transport (proton, carbohydrate, phosphate and protein import, protein insertion into membrane); protein modification (refolding, repair, targeting); chromosome (segregation, separation); biosynthesis (coenzyme A, dTMP, dTTP, lipoprotein); breakdown (deoxyribonucleotide, deoxyribose, mRNA, protein); regulation (phosphate, carbohydrate, and carboxylic acid metabolic processes, cellular phosphate ion homeostasis); cell-cell adhesion; foreign DNA cleavage; SOS response; sister chromatid cohesion; and uracil salvage.

These deletions reduce the ability of *M.genitalium* to interact with the environment and defend against external forces. This results in a reduction in control, from transport to regulation to genome management, and pruned metabolic processes and metabolites. This leaves Minesweeper_256’s *in-silico* cell alive, but more vulnerable to external and internal pressures, less capable of responding to change, and more reliant on internal processes occurring by chance.

In comparison, GAMA_236 has 163 genes (69% coverage) with GO terms on UniProt ^25^, with 73 genes with no GO terms. The 165 genes deleted reduced 17 (11%) GO categories, and removed 55 (35%) GO categories, 38 (69%) of which were associated with a single-gene (see Supplementary Information M).

8 unaffected and five reduced GO categories in Minesweeper_256 were removed in GAMA_236, with one unaffected GO category unique to GAMA_236 (phosphate ion transmembrane transport). Four GO categories were reduced further in GAMA_236: DNA (transcription, transcription regulation, transport) and glycerol metabolic process.

The 13 additional GO categories removed include: DNA (transcription (termination, regulation of elongation, antitermination, initiation)); RNA (processing (mRNA, tRNA, rRNA), rRNA catabolic process, tRNA modification, pseudouridine synthesis); thiamine (biosynthetic process, diphosphate biosynthetic process); and protein lipoylation.

GO analysis of GAMA_236, when compared to Minesweeper_256, suggests a further reduction of both internal control and reactivity to external environment.

### Genes with Low and High Essentiality

We analysed Minesweeper_256 and GAMA_236 to determine whether these were different minimal genomes, or GAMA_236 was an extension of Minesweeper_256. We conducted a gene content comparison of an unmodified *M.genitalium*, Minesweeper_256, and GAMA_236 genomes (Figure 4, Supplementary Information F), highlighting gene deletions unique to each minimal genome. We took this a step further and compared Minesweeper_256 to all of the GAMA genomes 256 to 236 genes in size. Figure 5 shows the GAMA algorithm’s avenue of gene reductions converging to a minimal genome, but Minesweeper_256 is not on the same path of convergence.

**Figure 4.**
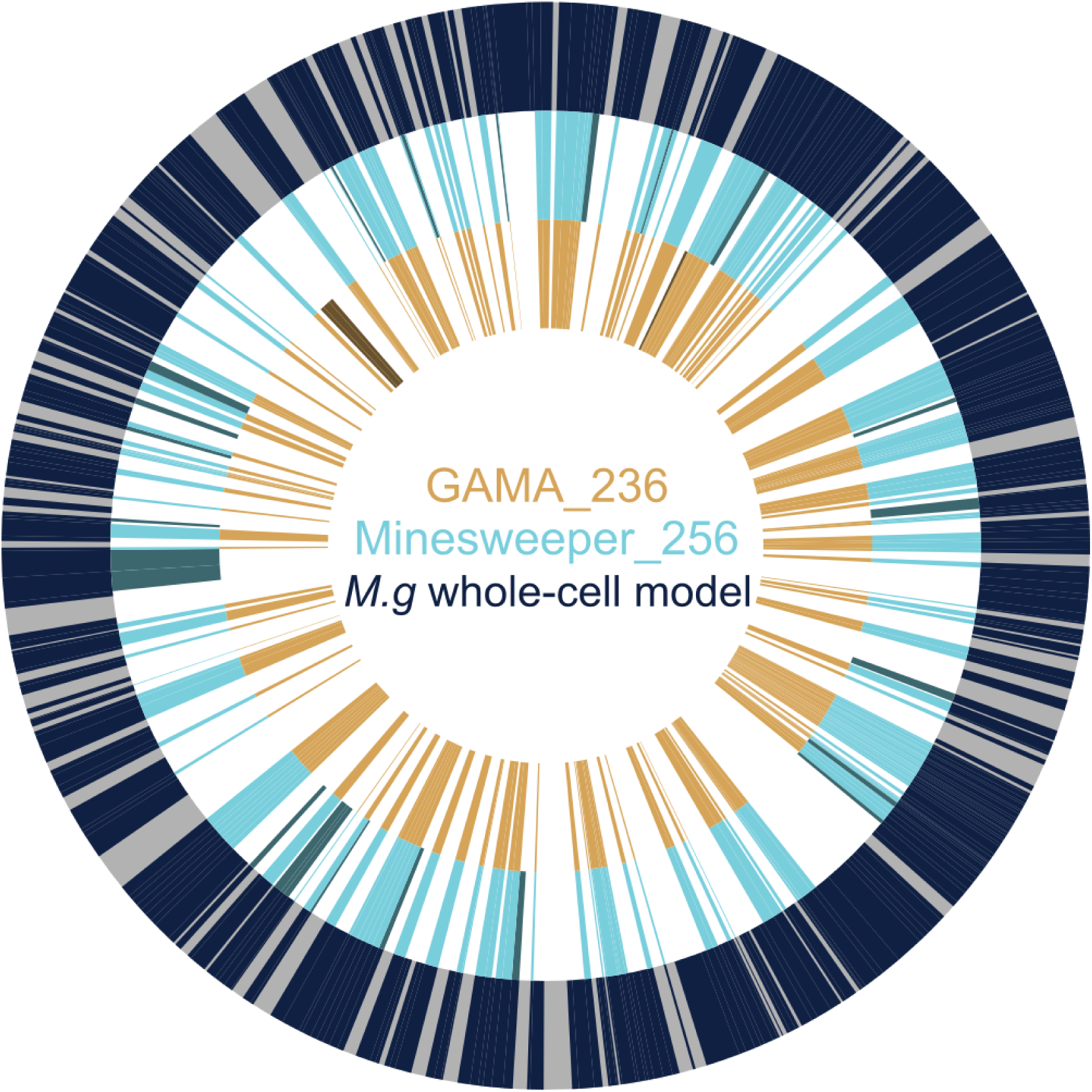
Comparing the genomes of the *Mycoplasma genitalium* whole-cell model, Minesweeper_256, and GAMA_236. The outer ring displays the *M.genitalium* genome (525 genes in total), with modelled genes (401) in navy and unmodelled genes (124, with unknown function) in grey. The middle ring displays the reduced Minesweeper_256 (256 genes) genome in light blue, with genes present in Minesweeper_265 but not in GAMA_236 in dark blue. The inner ring displays the reduced GAMA_236 (236 genes) genome in light yellow, with genes present in GAMA_236 but not in Minesweeper_265 in dark yellow. Figure produced from published *M.genitalium* genetic data ^17,18^, with genetic data for Minesweeper_256 and GAMA_236 available in the Supplementary Information.

**Figure 5.**
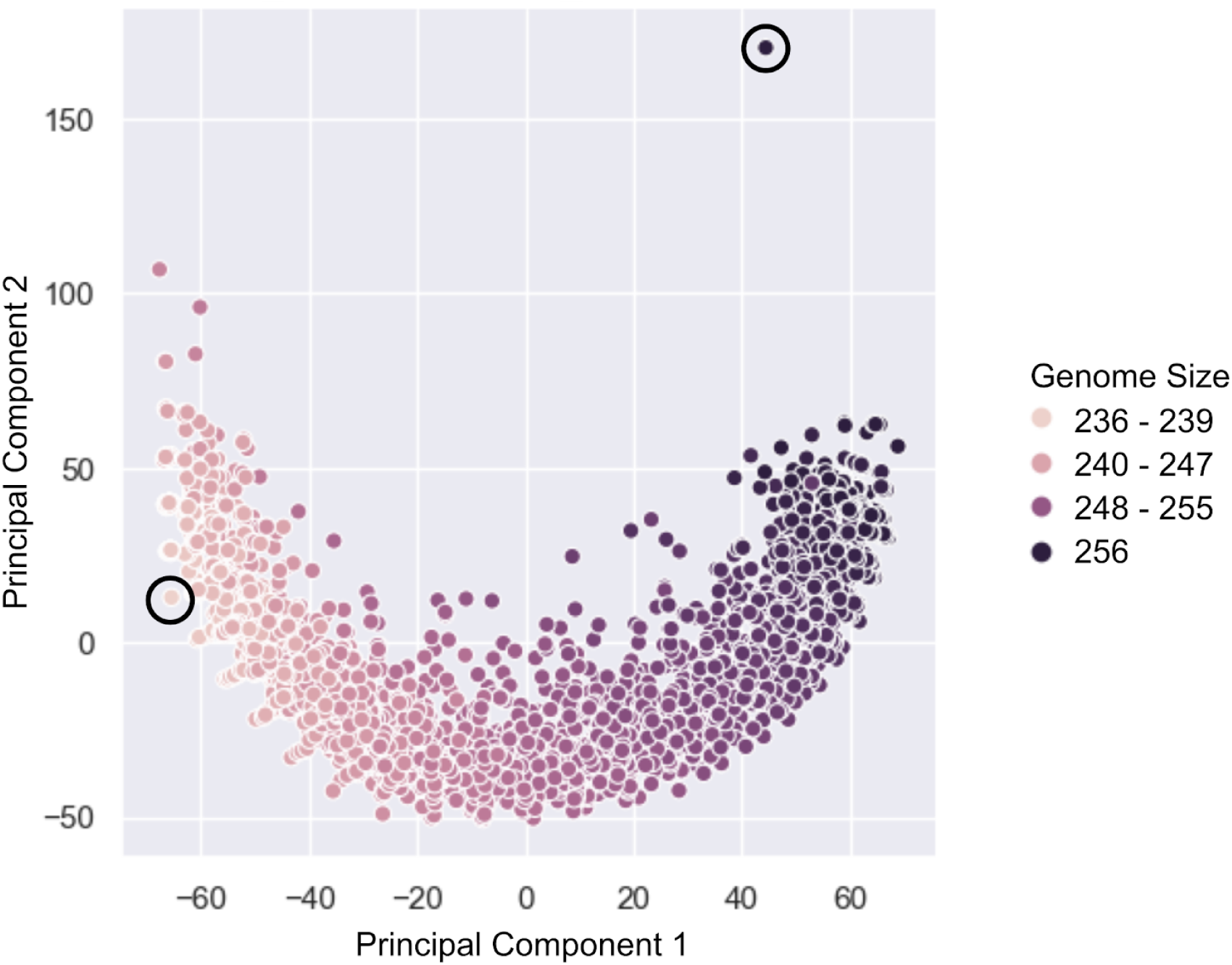
Comparing the genomes of Minesweeper_256 and 2954 GAMA genomes. The genome of Minesweeper_256 and all the genomes found by GAMA (that were the same size or smaller) were collated. Each point represents a single genome and is plotted based on a similarity metric (see Methods). The circled genome in the top right is Minesweeper_256 and the circled genome in the bottom left is GAMA_236. The key difference between the genomes is phosphate sources, with Minesweeper_256 using phosphonate and the GAMA genomes using inorganic phosphate.

Our comparison of the genomes found 18 genes knocked out in GAMA_236 that have high essentiality ^1^. They were defined as essential by single knockout in an unmodified *M.genitalium* whole-cell model, but could be removed in the genomic context of GAMA_236 without preventing division (see Supplementary Information A & E). We also found that four of these 18 genes could be removed as a group in the genomic context of Minesweeper_256, but doing so greatly increased the number of non-dividing cells produced (see Supplementary Information E).

Our genome comparison also found that Minesweeper_256 removed four genes, and GAMA_236 removed five genes (Table 1), which could not be removed either individually or as a group from its counterpart, without causing cellular death or mutations that prevented cellular division. We confirmed that these nine genes were individually non-essential. One additional gene, MG_305, could not be additionally removed in both GAMA_236 and Minesweeper_256. Our results demonstrate that these nine genes have low essentiality ^1^. To identify the cause of this synthetic lethality we attempted to match the functions of these low essentiality genes (Table 1), as we anticipated finding redundant essential gene pairs or groups. We found two genes in GAMA_236 (MG_289, MG_291) had matching GO terms with the gene MG_411 in Minesweeper_256. These, and three other adjacent genes on the genome, were tested by combinatorial gene knockouts in an unmodified *M.genitalium* whole-cell model genome (see Supplementary Information H). MG_289, MG_290, MG_291 were found to form a functional group, as were MG_410, MG_411, MG_412. These genes could be deleted individually and in functional groups from an otherwise unmodified *M.genitalium* whole-cell genome, and produce a dividing *in-silico* cell. However, any double gene deletion combination that involved one gene from each functional group resulted in a cell that could not produce RNA, produce protein, replicate DNA, grow or divide.

**Table 1.** Low Essentiality Genes from Minesweeper_256 and GAMA_236 genomic contexts. Protein annotation and GO term obtained from KEGG ^26^ and UniProt ^25^, based on Fraser et al’s *Mycoplasma genitalium\** G37 genome ^18^.

| Gene | Annotation | GO Term<br>(Biological Processes) | Non Essential In | Essential In |
| --- | --- | --- | --- | --- |
| MG_039 | N/A | N/A | GAMA_236 | Minesweeper_256 |
| MG_289 | p37 | transport | GAMA_236 | Minesweeper_256 |
| MG_290 | p29 | N/A | GAMA_236 | Minesweeper_256 |
| MG_291 | p69 | transport | GAMA_236 | Minesweeper_256 |
| MG_427 | N/A | OsmC-like protein | GAMA_236 | Minesweeper_256 |
| MG_033 | glpF | glycerol metabolic process | Minesweeper_256 | GAMA_236 |
| MG_410 | pstB | N/A | Minesweeper_256 | GAMA_236 |
| MG_411 | pstA | phosphate ion<br>transmembrane transport<br>process | Minesweeper_256 | GAMA_236 |
| MG_412 | N/A | N/A | Minesweeper_256 | GAMA_236 |
| MG_305 | dnaK | protein folding | *M.g* * whole-cell model | GAMA_236 and<br>Minesweeper_256 |

*M.genitalium* only has two external sources of phosphate, inorganic phosphate and phosphonate. MG_410, MG_411, and MG_412 transport inorganic phosphate into the cell, with MG_289, MG_290, and MG_291 transporting phosphonate into the cell ^18,26^. These phosphate sources proved to be a key difference between our minimal genomes. Minesweeper_256 removed the phosphate transport genes, relying on phosphonate as the sole phosphate source. GAMA_236 removed the phosphonate transport genes, relying on inorganic phosphate as the sole phosphate source. This can be seen in the GO term analysis, the phosphate ion transmembrane transport is still present in GAMA_236 but not in Minesweeper_256.

It has previously been theorised that individual bacterial species will have multiple minimal genomes ^27,28^, with different gene content depending on the environment and which evolutionary redundant cellular pathways were selected during reduction. We would argue that one of these selected pathways is phosphate source, with minimal genomes differing by choice of phosphate transport genes and associated processing stages, equivalent to the *phn* gene cluster in *Escherichia coli* ^*29*^. We could not however find any annotated phosphonate processing genes that had been subsequently removed in GAMA_236. We suspect that further “pivot points”, the selection of one redundant cellular pathway over another during reduction, will be identified in future *in-vivo* and *in-silico* bacterial reductions increasing the base number of minimal genomes per bacterial species.

## Discussion

We created two genome design algorithms (Minesweeper and GAMA) that used computational design-simulate-test cycles to produce *in-silico M.genitalium* minimal genomes (achieving 36% and 41% reductions, respectively). Our minimal genomes are smaller than *JCVI-syn3.0* (currently the smallest genome that can be grown in pure culture ^2^) and 33 - 53 genes smaller than the most recent predictions for a reduced *Mycoplasma* genome ^16^.

Additionally, we identified 10 low essentiality genes, 18 high essentiality genes ^1^, and produced evidence for at least two minima for Mycoplasma genitalium *in-silico*. We plan to test these results experimentally to ascertain the accuracy of the model and the functionality of our minimal genomes.

We believe that single-gene knockout classifications are unreliable for genome minimisation, as they fail to take into account genomic context. Single-gene knockout studies will underestimate minimal genome size as low essentiality genes will be scored as non-essential ^2,15,16^, but they will also overestimate minimal genome size as high essentiality genes will be scored as essential. We found 10 low essential genes within 358 protein-coding genes. As a single synthetic lethality event will prevent a genome from surviving, this gives a 3% chance of error for untested genome designs in even this evolutionarily reduced genome. Additionally, single-gene knockout studies narrow the scope of genome design; the 18 high essentiality genes identified as dispensable within GAMA_236 would not have been traditionally targeted by laboratory methods.

There are limitations to the approach presented here. Models are not perfect representations of reality: through necessity this model bases some of its parameters on data from other bacteria ^17^; multi-generation simulations are only possible by isolating one submodel from the rest of model (which loses genomic context); and *M.genitalium* has genes of unknown function that the model cannot account for.

The success of our *in-silico* genomes *in-vivo* is dependent on the accuracy of the model, which is untested at this scale of genetic modification. Minesweeper_256 and GAMA_236 may only function in the first generation of cells and the impact of the unmodelled genes is unknown. These genes may change the genomic context such that our minimal genomes are not successful, or as found with JCVI-Syn3.0 the genes of unknown function will be required for viability ^2^.

Our algorithms are currently adaptable to future, under development whole-cell models, as the algorithms interact with the models only via the input of gene deletion lists and analysing the output. This includes the *E.coli* whole-cell model at the Covert Lab, Stanford and the *Mycoplasma pneumoniae* whole-cell model at the Karr Lab, Mount Sinai, New York ^30^.

We believe that a hybrid of computational and lab based genome design and construction is now possible. This could produce quicker and cheaper laboratory results than currently possible, opening up this research to broader and interdisciplinary research communities. It also expands our research horizons raising the possibility of building truly designer cells, with increased efficiency and functional understanding.

## Methods

### Model Availability

The *M.genitalium* whole-cell model is freely available: https://github.com/CovertLab/WholeCell. The model requires a single CPU and can be run with 8GB of RAM. We run the *M.genitalium* whole-cell model on Bristol’s supercomputers using MATLAB R2013b, with the model’s standard settings. However, we use our own version of the SimulationRunner.m. MGGRunner.m is designed for use with supercomputers that start hundreds of simulations simultaneously, artificially incrementing the time-date value for each simulation, as this value is subsequently used to create the initial conditions of the simulation. This incrementation prevents the running of multiple simulations with identical initial conditions.

Our research copy of the whole-cell model was downloaded 2017-01-10.

### Code Availability

The code used for this research is openly available on Github (public code provided on publication). This includes the code for Minesweeper and GAMA genome design tools, scripts for statistical analysis, scripts for analysing GO terms, our custom simulation runner, analysis scripts, a template bash script, as well as the bash scripts and text files used to generate the simulations in this paper.

### Statistics

We used the R binom package (https://www.rdocumentation.org/packages/binom) to conduct one-tailed binomial proportion confidence intervals on our 41 genes showing inconsistent results (success ranging from 6 to 9 replicates, out of a total of 10 replicates). We used binom.confit.exact (Pearson-Klopper) using 95% CIs, producing for: 6/10 replicates [0.26, 0.87], 7/10 replicates [0.34, 0.93], 8/10 replicates [0.44, 0.97], 9/10 replicates[0.55, 0.99]). We graphed these results in R and in Python using Seaborn (https://seaborn.pydata.org/), the exact values, code, and graphs produced are available in Supplementary Information B & N.

Figure 5 was generated by creating a similarity matrix between all of the 2955 genomes, with the gene information represented in a binary format (present or absent). The matrix calculated a distance metric (1 - Adjusted Rand Index), with each genome comparison given a normalised score (0 = the genomes were identical, 1 = as different as would be expected if each genome was generated randomly, 2 = completely different). The resulting 2955 x 2955 matrix was then reduced to two dimensions with a standard PCA.

### Minesweeper

Minesweeper is written in Python3 and consists of four scripts (one for each stage). It uses no external libraries, so should be able to be run on any modern operating system (as they come with Python preinstalled) via a terminal. Each stage/script requires a text file(s) as input, with each stage outputting simulation files. These are run on a supercomputer and the automatically produced summary file is used as input for the next stage. Stages one to three are sequential, with stage four repeating until Minesweeper stops. Detailed instructions are provided in the README and progress is recorded in the deletion log in /OUTPUT_final.

The first stage of Minesweeper is optional, if you already have single gene knockout simulation results, you can proceed to the second stage. The second stage creates 26 deletion segments: 100%, 90%A, 90%B, 80%A, 80%B, 70%A, 70%B, 60%A, 60%B, 50%A, 50%B, 33%A-C, 25%A-D, 12.5%A-H. The A segments start from the top of the list of genes, whereas the B segments start from the bottom of the list of genes. The third stage progresses with the three largest deletion segments that produced a dividing cell, these three variants are referred to as red, yellow, blue. These perform as replicates and as a check on if the results are converging. The three variants are matched with smaller, dividing, non-overlapping segments using a list of allowed matches (implementation is detailed in third stage script), and unique combinations generated using a python implementation of powersets. The fourth stage splits the remaining genes into eight groups. The reason for selecting eight groups and three variants, is that a set of eight produces 256 unique combinations. Three variants each with 256 simulations (768 total) is 85% of the capacity of BlueGem. A set of nine groups with three variants (1536 simulations total) is 170% the capacity of BlueGem. Queueing systems mean that you don’t require this number of CPUs in total, but the execution time is multiplied as you wait for the simulations to process. The number of variants and groups can be lowered or increased depending on the number of CPUs you have available.

### GAMA

GAMA is written in Python3 and relies on a variety of different packages. These dependencies can be easily taken care of by installing it from PyPI using either ‘pip install genome_design_suite’ or ‘conda install genome_design_suite’ (it is recommended that you do this from within a virtual environment since this is pre-alpha and has not been extensively tested with different versions of all the libraries). A dependencies list is available in the main directory of the github repository if you would like to do this manually. The main dependency is the ‘genome_design_suite’ which is a suite of tools created by Oliver Chalkley at the University of Bristol which enables it to be easily run on different (or even multiple) clusters and well as enabling automatic data processing and database management. Due to the large amount of data produced by the Whole-Cell model, the simulation output data was reduced to essential data, converted into Pandas DataFrames (https://pandas.pydata.org/) and saved in Pickle files. GAMA would have produced 100s of TBs of data in the model’s native output format (compressed matlab files) which we are not able to store so this was an essential step. In order to run this code you must have a computer dedicated to remotely managing the simulations. A PC with a quad-core Intel(R) Xeon(R) CPU E5410 (2.33GHz) and 1GB of RAM running CentOS-6.6 was used as our computer manager, which is referred to as OC2. GAMA was run on OC2 using the scripts contained in gama_manamgement.zip Each stage of GAMA was run individually and manually updated as it was in proof-of-concept stage when GAMA_236 was found. ko.db is an SQLite3 database used to stored key information about simulations like average growth rate and division time.

The guess stage splits the singularly non-essential genes in roughly equally sized partitions. The four files, focus_on_NE_split_[1-4].py, run the exploration of each of the four partitions of the guess stage from OC2 - after unzipping gama_management.zip these can be found in gama/guess. The submission scripts and other files automatically created to run the simulations on the cluster can be found in gama_run_files.zip -> gama_run_files/guess. The simulation output is saved in Pickle files and can be found in gama_data/guess. Due to a technical problem the growth rate and division time of the genomes simulated in this stage are not in ko.db. viability_of_ne_focus_sets_pickles.zip contains the viability data of these simulations and the Python script used to collect it.

The add stage was executed on OC2 by running the files in gama_management.zip -> gama/add. The submission scripts and other files automatically created to run the simulations on the cluster can be found in gama_run_files.zip -> gama_run_files/add. The simulation output can be found in gama_data/add and an overview of the simulation results can be found in ko.db where the batchDescrription.name is some derivative of ‘mix_ne_focus_split’.

The mate stage was executed on OC2 by running the file in gama_management.zip -> gama/mate. The submission scripts and other files automatically created to run the simulations on the cluster can be found in gama_run_files.zip -> gama_run_files/mate. The simulation output can be found in gama_data/mate and an overview of the simulation results can be found in ko.db where batchDescription.name is some derivative of ‘big_mix_of_split_mixes’.

### Equipment

We used the University of Bristol Advanced Computing Research Centres’s BlueGem, a 900-core supercomputer, which uses the Slurm queuing system, to run whole-cell model simulations. GAMA also used BlueCrystal, a 3568-core supercomputer, which uses the PBS queuing system. We used a standard office desktop computer, with 8GB of ram, to write new code, interact with the supercomputer, and run single whole-cell model simulations. We used the following GUI software on Windows/Linux Cent OS: Notepad++ for code editing, Putty (ssh software)/the terminal to access the supercomputer, and FileZilla (ftp software) to move files in bulk to and from the supercomputer. The command line software we used included: VIM for code editing, and SSH, Rsync, and Bash for communication and file transfer with the supercomputers.

### Data Format

The majority of output files are state-NNN.mat files, which are logs of the simulation split into 100-second segments. The data within a state-NNN.mat file is organised into 16 cell variables, each containing a number of sub-variables. These are typically arranged as 3-dimensional matrices or time series, which are flattened to conduct analysis. The other file types contain summaries of data spanning the simulation.

### Data Analysis Process

The raw data is automatically processed as the simulation ends. runGraphs.m carries out the initial analysis, while compareGraphs.m overlays the output on collated graphs of 200 unmodified *M.genitalium* simulations. Both outputs are saved as MATLAB .fig and .pdfs, though the .fig files were the sole files analysed. The raw .mat files were stored in case further investigation was required. To classify our data we chose to use the phenotype classification previously outlined by Karr (Figure 6B ^17^), which graphed five variables to determine the simulated cells’ phenotype. However, the script responsible for producing Figure 6B, SingleGeneDeletions.m, was not easily modified. This led us to develop our own analysis script recreating the classification: runGraphs.m graphs growth, protein weight, RNA weight, DNA replication, cell division, ands records several experimental details. There are seven possible phenotypes caused by knocking out genes in the simulation: non-essential if producing a dividing cell; and essential if producing a non-dividing cell because of a DNA replication mutation, RNA production mutation, protein production mutation, metabolic mutation, division mutation, or slow growing.

For the single gene knockout simulations produced in initial input, the non-essential simulations were automatically classified and the essential simulations flagged. Each simulation was investigated manually and given a phenotype manually using the decision tree (see Supplementary Information D). For simulations conducted by Minesweeper and GAMA, simulations were automatically classified solely by division, which can be analysed from cell width or the endtime of the simulation.

Further analysis, including: cross-comparison of single-gene knockout simulations, comparison to Karr et al’s ^17^ results, analysis of Minesweeper and GAMA genomes (genetic content and similarity, behavioural analysis, phenotypic penetrance, gene ontology), and identification and investigation of high and low essentiality genes and groupings, were completed manually. The GO term analysis of gene deletion impacts was processed by a created script (see Github for code), then organised into tables of GO terms that were unaffected, reduced, or removed entirely.

### Modelling: Scripts, Process and Simulations

Generally, there are six scripts we used to run the whole-cell model. Three are the experimental files created with each new experiment (the bash script, gene list, experiment list), and three are stored within the whole-cell model and are updated only upon improvement (MGGrunner.m, runGraphs.m, and compareGraphs.m). The bash script is a list of commands for the supercomputer(s) to carry out. Each new bash script is created from the GenericScript.sh template, which determines how many simulations to run, where to store the output, which analysis to run, and where to store the results of the analysis. The gene list is a text file containing rows of gene codes (in the format ‘MG_XXX’,). Each row corresponds to a single simulation and determines which genes that simulation should knockout. The experiment list is a text file containing rows of simulation names. Each row corresponds to a single simulation and determines where the simulation output and results of the analysis are stored. In brief, to manually run the whole-cell model: a new bash script, gene list, and experiment list are created on the desktop computer to answer an experimental question. The supercomputer is accessed on the desktop via ftp software, where the new experimental files are uploaded, the planned output folders are created, and MGGRunner.m, runGraphs.m, compareGraphs.m files are confirmed to be present. The supercomputer is then accessed on the desktop via ssh software, where the new bash script is made executable and added to the supercomputer’s queuing system to be executed. Once the experiment is complete, the supercomputer is accessed on the desktop via ssh software, where the results of the analysis are moved to /pdf and /fig folders. These folders are accessed on the desktop via ftp software, where the results of the analysis are downloaded. More detailed instructions are contained within the template bash script.

Each wild-type simulation consists of 300 files requiring 0.3GB. Each gene manipulated simulation can consist of up to 500 files requiring between 0.4GB and 0.9GB. Each simulation takes 5 to 12 hours to complete in real time, 7 - 13.89 hours in simulated time.

### Data Availability

The databases used to design our *in-silico* experiments, and compare our results to, includes Karr et al ^17^ and Glass et al ^24^ Supplementary Information, and Fraser et al *M.genitalium* G37 genome ^18^ interpreted by KEGG ^26^ and UniProt ^25^ as strain ATCC 33530/NCTC 10195.

Minesweeper simulations raw and transformed output (.mat files) are available upon request, as the they require 4.2 TB of storage. The output .fig files (10 GB) are available for download from the our group’s Research Data Repository at the University of Bristol. GAMA simulations transformed output is available in ko.db.

## Acknowledgments

We would like to thank the Advanced Computing Research Centre (ACRC) and BrisSynBio, a BBSRC/EPSRC Synthetic Biology Research Centre, at the University of Bristol for access to the BlueCrystal and Bluegem supercomputers. Special thanks to the HPC and RDSF teams of the ACRC, particularly Dr. Christopher Woods, Simon Burbidge, Matt Williams, and Damian Steer for their help with BlueCrystal, BlueGem, data storage and publication.

We would like to thank Jonathan Karr for his advice on running gene knockout simulations using the *M. genitalium* whole-cell model, and for his constructive and enlightening feedback.

We would like to thank Anthony Vecchiarelli (Assistant Professor, University of Michigan) and class MCDB 401 (“Building the Synthetic Cell”) for conducting a class review of our preprint paper, providing us with constructive and encouraging feedback.

We would like to thank John Glass (Professor, JCVI Synthetic Biology and Bioenergy Group) for his constructive and informative feedback.

We would like to thank Cameron Matthews and Julia Needham, University of Bristol undergraduates, who conducted simulations to test the inclusion of MG_290 within the phosphonate group as part of summer studentships.

LM is supported by the Medical Research Council grant MR/N021444/1 to LM, and by the Engineering and Physical Sciences Research Council grant EP/R041695/1 to LM.

OC, LM and CG are supported by a BrisSynBio, a BBSRC/EPSRC Synthetic Biology Research Centre (BB/L01386X/1), flexi-fund grant.

OC is supported by the Bristol Centre for Complexity Sciences (BCCS) Centre for Doctoral Training (CDT) EP/I013717/1; JR and SL are supported by EPSRC Future Opportunity Scholarships.

## Author Contributions

C.G, L.M, O.C for attaining initial funding.

C.G, L.M, O.P, O.C, J.R, S.L were involved in ideation.

S.L was involved in analysis and development of Figure 4.

O.C was responsible for the development and implementation of the *Mycoplasma genitalium* whole-cell model outside of the Covert Lab, Stanford (on Bristol’s BlueGem and BlueCrystal), initial ideation about uses of whole-cell models, GAMA (method, results, section), Figure 2, Figure 5, and collaborative theorising on essentiality and minimal genomes.

J.R was responsible for the development of automated graphing, Minesweeper (method, results), spreadsheet analysis of *in-silico* results, Table 1, Figure 1, Figure 3, collaborative theorising on essentiality and minima, and writing of the paper and Supplementary Information.

C.G, L.M, O.C, O.P, S.L were involved in editing and feedback on paper.

## Competing Interests

The authors declare no competing interests.

